# Autism-related behavioral phenotypes of three *Camk2a* mutant mouse lines with reduced CaMKIIα expression and/or activity

**DOI:** 10.1101/2022.10.28.514232

**Authors:** Jessica R. Thomas, Keeley L. Spiess, Roger J. Colbran

## Abstract

Ca^2+^/calmodulin-dependent protein kinase II (CaMKII) is a key modulator of excitatory synaptic transmission, gene expression, learning and memory. Mutations in the *CAMK2A* gene, which encodes CaMKIIα and is highly expressed in multiple regions in the forebrain, have been recently linked to neurodevelopmental disorders such as autism spectrum disorder (ASD) and intellectual disability (ID). Our lab generated and characterized a knock-in (KI) mutant mouse with a glutamate-183 to valine (E183V) CaMKIIα mutation detected in several children diagnosed with ASD or ID. The E183V mutation reduces CaMKIIα activity and expression levels but the contributions of these two changes to the ASD-related behavioral phenotypes of these mice are unclear. Therefore, we performed side-by-side comparisons of the behavioral phenotypes of CaMKIIα E183V-KI mice with two other mutant mouse lines with either a complete loss of CaMKIIα expression (CaMKIIα Null mice) or reduced kinase activity (due to a threonine-286 to alanine mutation that abrogates autophosphorylation at this site) with no significant change in expression levels (CaMKIIα T286A-KI mice). In all three lines, homozygous mutant mice displayed increased stereotypic jumping behavior and hyperactivity, without alterations in anxiety or social interactions. Interestingly, homozygous mutant mice in all three lines also displayed a substantial reduction in tactile sensitivity using the Von Frey filament test. Together, these data suggest that reductions of either CaMKIIα expression or activity in mice disrupted normal motor and sensory functions.

## Introduction

Ca^2+^/calmodulin-dependent protein kinase (CaMKII) is a serine/threonine kinase that is encoded by four mammalian genes (*CAMK2A, CAMK2B, CAMK2G, CAMK2D* in humans) ^1-3^. A single CaMKII subunit contains catalytic, regulatory and association domains. In the basal state, the regulatory domain interacts with the catalytic domain to block the active site and suppress kinase activity ^4,5^. This interaction is disrupted by Ca^2+^/calmodulin binding to the regulatory domain, exposing the ATP- and substrate-binding sites to activate the kinase ^6-8^. Self-oligomerization of the association domain forms the dodecameric hub of a holoenzyme, facilitating efficient autophosphorylation between adjacent activated subunits in the regulatory domain (at Thr286 in CaMKIIα; Thr287 in other isoforms) ^9-12^. Thr286/287 autophosphorylation prevents the regulatory domain from interacting with the catalytic domain following Ca^2+^/calmodulin dissociation, resulting in autonomous (Ca^2+^-independent) CaMKII activity ^13-15^.

Given abundant evidence that CaMKII is a critical regulator of the synaptic processes that underlie learning and memory ^16-18^, it is perhaps surprising that mutations in the *CAMK2A, CAMK2B* and *CAMK2G* genes were only recently linked to neurodevelopmental and neurological disorders, including autism spectrum disorder (ASD), intellectual disability (ID) and epilepsy ^19-22^. Non-synonymous mutations in the catalytic, regulatory, and association domains have been functionally characterized to varying extents ^19,21-26^. Our lab characterized a unique ASD-linked *de novo CAMK2A* mutation, encoding a glutamate-183 to valine (E183V) substitution in the catalytic domain of the CaMKIIα isoform, which is abundantly expressed in many parts of the forebrain. This same mutation was later identified in other children with ID or ASD ^20,26^. *In vitro* studies showed that the E183V mutation severely reduced CaMKIIα kinase activity, Thr286 autophosphorylation and binding to multiple CaMKII-associated proteins (CaMKAPs). Expression of E183V-CaMKIIα in cultured hippocampal neurons disrupted dendritic outgrowth and dendritic spine formation, and reduced excitatory synaptic transmission. Moreover, homozygous mice with the CaMKIIα E183V knock-in mutation (E183V-KI mice) displayed ASD-related behavioral phenotypes: impaired social interactions, repetitive behaviors, and hyperactivity. However, in addition to the loss of kinase activity and Thr286 autophosphorylation, CaMKIIα expression was substantially reduced in E183V-KI mice. As a result, ASD-related behaviors in CaMKIIα E183V-KI mice may be due to reductions of CaMKIIα expression and/or activity.

While reduction or loss of CaMKIIα expression and/or activity in mice generally alters synaptic plasticity and behavior, some differences in phenotypes have been reported. For example, loss of CaMKIIα expression resulted in enhanced short-term plasticity, while complete or partial loss of CaMKIIα activity did not impact short-term plasticity ^27-29^. Moreover, loss of CaMKIIα expression or activity impaired spatial memory, but visually guided memory remained intact when CaMKIIα activity, but not expression, was abolished ^16,30^. In addition, the functional impact of losing CaMKIIα expression or activity may be brain region specific ^30,31^. For example, hippocampal-dependent memory, such as contextual memory, appears to require CaMKIIα activity, while amygdala-dependent cued fear memory may not ^30^. However, some of the variability in these reported phenotypes may arise from differences in the strain background and/or housing conditions of *Camk2a* mutant mice tested. These issues can only be resolved by making side-by-side comparisons of the *Camk2a* mutant mice.

Here, we report side-by-side phenotypic comparisons of CaMKIIα Null mice that lack CaMKIIα expression, CaMKIIα T286A-KI mice with reduced autonomous CaMKIIα activity due to a lack of Thr286 autophosphorylation, and CaMKIIα E183V-KI mice with reduced CaMKIIα expression and CaMKIIα activity. We compare social, motor, and sensory behaviors, hyperactivity and anxiety. We found that all three mouse lines have similar impairments in motor behaviors and sensory functions, but no substantial impairments in social functions and anxiety. Moreover, the paws of all three mouse lines displayed robust tactile hyposensitivity in the Von Frey filament test.

## Materials and Methods

### Animals

Mice (2-5 mice) were housed in standard cages with paper bedding on a 12-hour light/dark cycle (light: 6:00 A.M. to 6:00 P.M.) with food and water ad libitum. CaMKIIα E183V-KI mice (MGI: 5811610) were described previously ^26^, and were backcrossed at least six times to a C57BL/6J (B6) background for the current studies. CaMKIIα Null mice also were previously described ^32^ and backcrossed to a B6 background. CaMKIIα T286A-KI mice (MGI:2158733) ^33^ also were on a B6 background. Genotyping was performed before and after behavioral testing. Genotyping assays for CaMKIIα Null and T286A mice were previously described ^32,34^. Primers used to genotype the CaMKIIα E183V-KI mice are as follows:

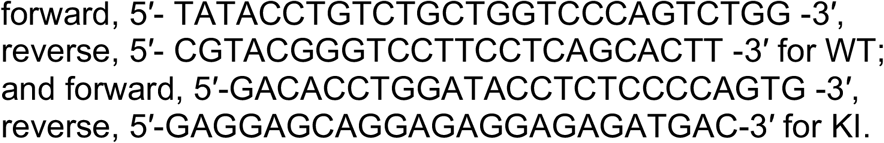

We tested 2-3 cohorts of the three CaMKIIα mutant mouse lines, with each cohort containing the random mixture of male and female mice of each genotype generated using a standard HETXHET breeding strategy. Behavioral tests were performed between the ages of 7 and 18 weeks of age in the order listed below, with the exception that some data included in figures 3 (3-chamber social test) and 7 (Von Frey filament test) were obtained using naïve mice, as described below. The experimenter was blinded to genotypes during testing.

All mouse experiments were approved by the Vanderbilt University Institutional Animal Care and Use Committee and were performed in accordance with the National Institutes of Health Guide for the care and use of laboratory animals.

### *Novel open-field locomotor activity* (7-14 weeks)

Naïve mice were placed in the center of a 27 × 27 cm open-field chamber housed in sound-attenuating case (Med Associates) for a 60-min. period. The center zone of the open-field box was pre-defined (19.05 × 19.05 cm) in the Activity Monitor software (Med Associates; RRID:SCR_014296), which recorded infrared beams breaks as movements in the center and outer zones in the x-, y-, and z-dimensions.

### *Elevated plus maze* (7-16 weeks)

The four arms of the maze were approximately 5-cm wide and elevated approx. 2 feet from the floor. The open arms had a 0.5 cm lip to prevent the mouse from falling whereas the walls of closed arms were 20-30 cm tall. Mice were placed at the intersection of the arms and permitted to explore freely for 10 min while being videotaped from above. Mouse behavior was tracked and measured using ANY-maze software (Stoelting; RRID:SCR_014289). The position of the mouse was scored based on the location of the head and trunk of the test mouse in the open arms, closed arms, or center of the maze (Stoelting).

### *Light-Dark box* (8-18 weeks)

The open-field chambers described above were divided into light and dark zones using a black Plexiglass insert measuring 13.5 × 27 cm. The test was started by placing mice in the light side of the chamber and giving them 10 mins to explore the light and dark zones of the box, as pre-defined using the Activity Monitor software (Med Associates). In addition, the software recorded infrared beams breaks in the x-, y-, and z-dimensions as movements in the pre-defined zones.

### *Three-chambered test* (10-18 weeks)

Social exploration was measure in a clear polycarbonate apparatus, covered with white paper towel to block visual cues, with 4-inch sliding gates separating three 7 × 9-inch chambers. Both side chambers contained an inverted metal wire pencil cup in one corner. There were two phases to the testing: (1) A 10 min habituation session allowed the test mouse to freely explore all three chambers with empty wire cups in both positions. (2) After guiding the test mouse into the center chamber, a WT stranger mouse of the same sex was placed under one of the two metal cups. The test mouse was then allowed to freely explore all three chambers for 10 min. Mouse behavior was videotaped from above and measured using ANY-maze software (Stoelting). Interactions with the stranger mouse cup and the empty wire cup were scored as the amount of time the head and trunk of the test mouse was located within 2 cm of the appropriate wire cup. Mice from an initial cohort were excluded from the final analysis due to a technical error with the choice of stranger mice, and additional naïve mice were tested; there was no obvious difference in phenotypes between naïve mice and mice that experienced the prior behavioral tests, so the data were pooled for inclusion in the final Figure 3.

### *Von Frey filament test* (7-13 weeks)

Mice were acclimated to a 10 × 14 cm chamber with a wire floor for 1 hr. Filaments of various diameters were pressed against the plantar surface of the foot, bending to produce a constant application force from 0.01 to 10 mN. Filaments were tested in ascending order until a foot withdrawal response was observed. This process was repeated three times on alternating hind paws (twice on one paw and once on contralateral hind pawpaw). For each mouse, the average of the three readings was calculated to determine the average force required to evoke a paw withdrawal. Additional data obtained using naïve mice were included in the final figure 7 because there was no obvious difference in phenotypes compared with mice that experienced the prior behavioral tests.

### Data analysis and statistics

A ROUT test was used to identify outliers. The D’Agostino & Pearson and Shapiro-Wilk tests were used to test for normality. Statistical analyses of normally-distributed data used a paired t-test, one-way analysis of variance (ANOVA), or repeated measures two-way ANOVA (GraphPad Prism 9.2.0, LaJolla, CA, USA; RRID:SCR_002798), as described in the figure legends. A Kruskal-Wallis test was used to test data that was not normally distributed. If ANOVAs revealed significant variation between groups, Bonferroni, Dunn’s, or Tukey *post hoc* test was performed for pair-wise comparisons of multiple groups, as recommended in Prism (see figure legends). P values of < 0.05 were considered statistically significant. The results of selected statistical comparisons are cited in the text or figure legends and complete Prism outputs from all statistical tests are reported in supplementary table 1. All data are plotted as the mean ± SEM with super-imposed values for individual mice shown in a scatter plot.

## Results

### Stereotypic phenotypes

CaMKIIα E183V-KI homozygous (HOM) mice on a mixed background exhibited sustained and robust increases in jumping and rearing behaviors when monitored for 30 min in the open field test ^26^. Therefore, we compared jumping behaviors of wild-type (WT), heterozygous (HET) or HOM mice from the three CaMKIIα mutant mouse lines using the open-field test. HOM CaMKIIα E183V-KI (E183V-KI) mice on the pure B6 background displayed an enhanced jumping phenotype over the total 60 min test period (**Fig 1A**). Likewise, HOM CaMKIIα Null and CaMKIIα T286A-KI (Null and T286A-KI, respectively) mice exhibited increased total jumping compared to WT littermates (**Fig 1B-C**). More detailed comparison of jumping behavior in 5 min time blocks revealed that the jumping behavior of WT and HET mice from all three lines progressively decreased during the 60 min test. HOM mice from all three lines displayed increased jumping during the first 5-10 min of the test, but the jumping behaviors of HOM E183V-KI and Null mice were more persistent and remained elevated compared to WT and HET littermates throughout the test. In contrast, jumping behavior of HOM T286A-KI decreased over time, and was not significantly different from WT or HET littermates during the last 10 min of the test (**Fig 1D-F**). These data suggest a reduction in CaMKIIα expression or activity is sufficient to induce a repetitive, stereotypic jumping behavior, where a complete loss of both activity and expression results in a somewhat more severe phenotype.

**Figure 1.**
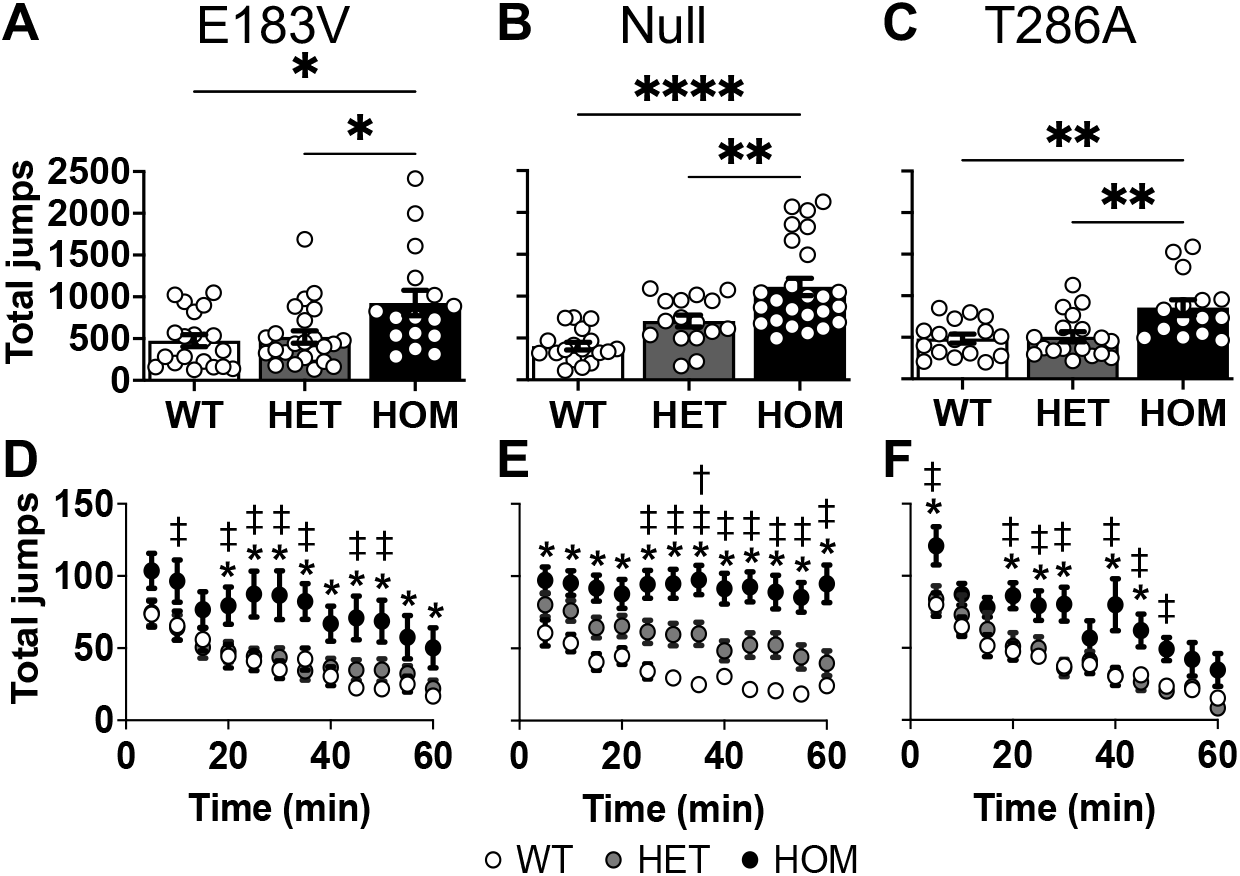
Stereotypic jumping during open-field testing of the three *Camk2a* mutant mouse lines. **A-C:** Total jumps in 60 min (mean ± SEM). E183V-KI (**A**): Kruskal-Wallis, P=0.012; Dunn’s *post hoc* analysis. Null (**B**): 1-way ANOVA, P<0.0001; Bonferroni’s *post hoc* analysis. T286A-KI (**C**): 1-way ANOVA, P=0.0008; Bonferroni’s *post hoc* analysis. Symbols for panels A-C indicate P ≤ 0.05 for *, P ≤ 0.01 for **, and P ≤ 0.001 for ***. **D-F:** Number of jumps in 5 min time windows (mean ± SEM) across the 60 min open-field test. All data analyzed with a 2-way repeated-measures ANOVA and Tukey’s *post hoc analysis* E183V-KI (**D**): interaction, F (22, 627) = 1.347, P=0.1337; genotype, F (2, 57) = 5.919, P=0.0046; time, F (11, 627) = 33.39, P<0.0001. Null (**E**): interaction, F (22, 616) = 1.962, P=0.0056; genotype, F (2, 56) = 18.15, p<0.0001; time, F (11, 616) = 7.966, P<0.0001. T286A-KI (**F**): interaction, F (22, 506) = 1.069, P=0.3768; genotype, F (2, 46) = 8.400, P=0.0008; time, F (11, 506) = 32.69, P<0.0001 Symbols for individual time points in panels D-F indicate P < 0.05 for: †, WT vs HET; *, WT vs HOM; ‡, HET vs HOM. Numbers of WT, HET and HOM mice, respectively, were: E183V-KI: 20, 24, 16. Null: 18, 16, 25. T286A-KI: 16, 18, 15.

### Hyperactivity

We next assessed the total distance traveled by each mouse line in the novel open-field test. HOM mice from all three mutant lines covered a significantly greater total distance over the 60 min test period relative to WT and HET littermates (**Fig 2A-C**). More detailed analysis of activity across the test period revealed that distances travelled by HOM mice in each line were significantly higher than distances traveled by HET and WT littermates during the initial 5-10 min of the test. Moreover, distances travelled by the three genotypes in each mouse line progressively decreased during each successive 5 min period, as expected due to habituation of the mice to the chambers. However, HOM CaMKIIα Null remained significantly hyperactive relative to WT and HET littermates throughout the 60 min test, whereas HOM E183V-KI and HOM T286-KI mice were significantly hyperactive relative to WT and HET littermates mostly during the first 20-30 min (**Fig 2D-F**).Taken together, these data suggest that decreasing CaMKIIα expression or activity is sufficient to induce hyperactivity, but that hyperactivity is somewhat more pronounced and persistent when both expression and activity are completely lost.

**Figure 2.**
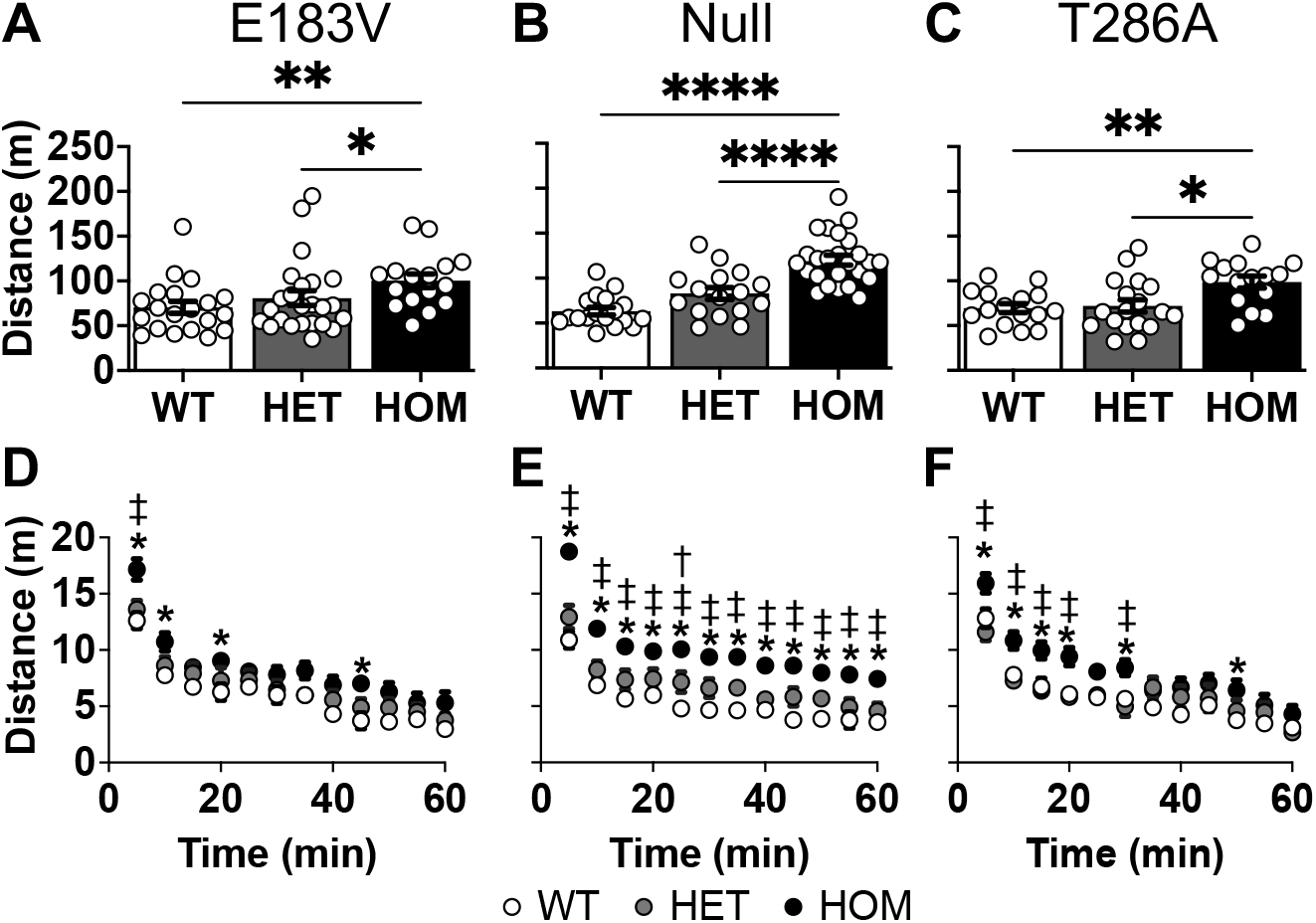
Distances travelled during open-field testing of the three *Camk2a* mutant mouse lines. **A-C**: Total distance traveled in 60 min (mean ± SEM). E183V-KI (**A**): Kruskal-Wallis P=0.0074; Dunn’s *post hoc* analysis. Null (**B**): 1-way ANOVA P<0.0001; Bonferroni’s *post hoc* analysis T286A-KI (**C**): 1-way ANOVA P=0.0043; Bonferroni’s *post hoc* analysis. Symbols panels A-C indicate P≤0.05 for *, P≤0.01 for **, and P≤0.0001 for ****. **D-F**: Distances traveled in 5 min time windows (mean ± SEM). All data analyzed using a 2-way repeated-measures ANOVA with Tukey’s *post hoc* analysis. E183V-KI (**D**): interaction, F (22, 627) = 1.178, P=0.2608; genotype, F (2, 57) = 3.342, P=0.0424; time, F (11, 627) = 110.1, P<0.0001. Null (**E**): interaction, F (22, 616) = 1.867, P=0.0097; genotype, F (2, 56) = 29.87, P<0.0001; time, F (11, 616) = 77.11, P<0.0001. T286A-KI (**F**): interaction, F (22, 506) = 2.025, P=0.0040; genotype, F (2, 46) = 6.146, P=0.0043; time, F (11, 506) = 66.29, P<0.0001. Symbols for individual time points in panels D-F indicate P < 0.05 for: †, WT vs HET; *, WT vs HOM; ‡, HET vs HOM. Numbers of WT, HET and HOM mice, respectively, were: E183V-KI: 20, 24, 16. Null: 18, 16, 25. T286A-KI: 16, 18, 15.

### Social exploratory behavior

Next, we assessed social interaction using the three-chamber social test (3CST), which is widely used to characterize social phenotypes for ASD mouse models ^35,36^. During the habituation phase, none of the mice distinguished between two empty wire cups (**Fig 3A-C**). During the test phase, WT, HET, and HOM E183V-KI mice spent significantly more time exploring the wire cup containing the mouse as opposed to the empty cup (**Fig 3D**). Similarly, WT and HET Null and T286A-KI mice spent significantly more time exploring the wire cup containing the mouse as opposed to the empty cup. In contrast, there was not a significant difference in the time that HOM Null and T286A-KI mice spent exploring the two cups (**Fig 3E-F**). Interestingly, a significant increase in the total distance traveled was observed for HOM E183V-KI, Null, and T286A-KI mice during the test phase compared to WT and HET littermates (**Fig 3G-I**), but there was no difference in the number of entries into the chambers containing an empty wire cup or a wire cup with a mouse underneath for WT, HET, or HOM amongst the three mouse lines (data not shown). These data confirm the hyperactivity of all three CaMKIIα mouse lines that was observed in the open-field test, and indicate that this hyperactivity correlates with a modest reduction in social exploration, at least in the CaMKIIα NULL and T286A-KI mice. Thus, reductions of CaMKIIα expression and/or activity has a modest impact on social exploratory behavior.

**Figure 3.**
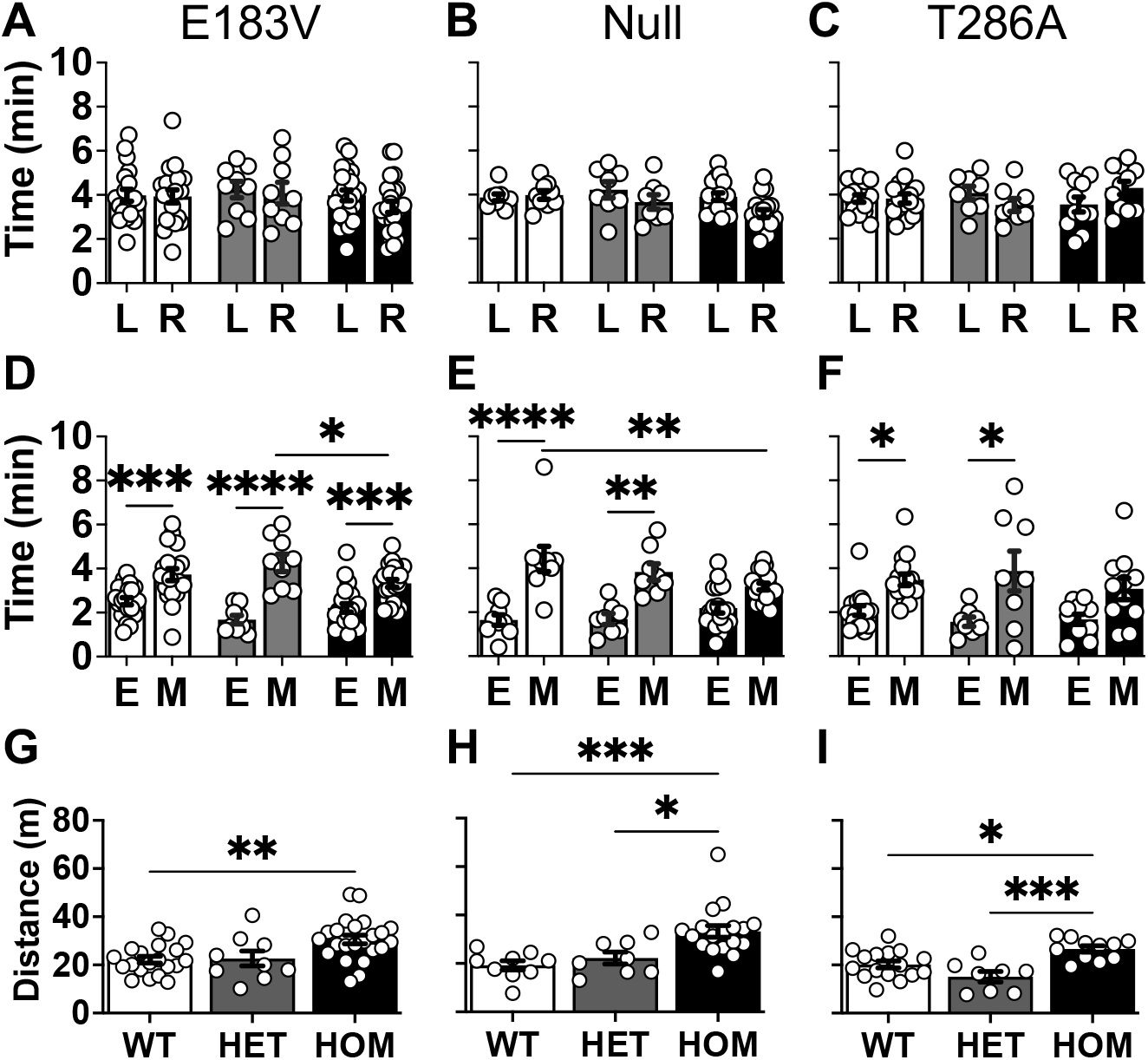
Social exploration by the three *Camk2a* mutant mouse lines in the 3CST. **A-C**. Times (mean ± SEM) spent exploring the left (L) versus the right(R) chamber for CaMKIIα E183V-KI (**A**), Null (**B**), and T286A-KI (**C**) mice. Panels **A-F** analyzed using a 2-way repeated-measures ANOVA with Bonferroni’s *post hoc* analysis. E183V-KI (**A**): interaction, F (2, 49) = 0.2152, P=0.8072; genotype, F (2, 49) = 7.258, P=0.0017; chamber, F (1, 49) = 0.4556, P=0.5029. Null (**B**): interaction, F (2, 32) = 1.093, P=0.3474; genotype, F (2, 32) = 14.58, P<0.0001; chamber, F (1, 32) = 2.379, P=0.1328. T286A-KI (**C**): interaction, F (2, 32) = 1.437, P=0.2525; genotype, F (2, 32) = 0.5982, P=0.5558; chamber, F (1, 32) = 0.05761, P=0.8118. **D-F**. Times (mean ± SEM) spent closely interacting with an empty wire pencil cup (E) versus an identical cup sheltering a novel mouse (M) for CaMKIIα E183V-KI (**D**), Null (**E**), and T286A-KI (**F**) mice. E183V-KI (**D**): interaction, F (2, 49) = 4.220, P=0.0204; genotype, F (2, 49) = 1.593, P=0.2136; chamber, F (1, 49) = 65.58, P<0.0001. Null (**E**): interaction, F (2, 32) = 3.744, P=0.0346 ; genotype, F (2, 32) = 1.481, P=0.2427; chamber, F (1, 32) = 41.89, P<0.0001. T286A-KI (**F**): interaction, F (2, 32) = 0.5436, P=0.5859; genotype, F (2, 32) = 1.151, P=0.3290; chamber, F (1, 32) = 19.63, P=0.0001. **G-I**. Distance traveled (mean ± SEM) for CaMKIIα E183V-KI (**G**), Null (**H**), and T286A-KI (**I**) mice. E183V-KI (**A**): Kruskal-Wallis, P=0.0025; Dunn’s *post hoc* analysis. Null (**B**): Kruskal-Wallis, P=0.0004; Dunn’s *post hoc* analysis. T286A-KI (**C**): 1-way ANOVA, F (2, 32) = 11, P=0.0003; Bonferroni’s *post hoc* analysis Symbols indicate P≤0.05 for *, P≤0.01 for **, P≤0.001 for ***and P≤0.0001 for ****. Numbers of WT, HET and HOM mice, respectively, were: E183V-KI: 20, 9, 23. Null: 9, 8, 18. T286A-KI: 16, 8, 11.

### Anxiety-related phenotypes

As an initial assessment of potential anxiety phenotypes in CaMKIIα mutant mice, we measured the amount of time spent in the center of the novel open-field arena. During the first 10 mins, HET and HOM E183V-KI mice spent less time in the center of the arena compared to WT littermates (**Fig 4A**), which is indicative of increased anxiety. HOM Null and T286A-KI mice, but not their HET littermates, spent less time in the center compared to WT mice (**Fig 4B-C**). However, the phenotypes appeared to change as the mice habituated to the open field arena. During the last 10 min in the open field arena, HET and HOM E183V-KI mice spent a similar amount of time in the center as their WT littermates (**Fig 4D**). Similarly, HOM Null mice spend an equal amount time in the center as their WT littermates, whereas HET Null mice spent more time in the center compared to WT and HOM littermates (**Fig 4E**), indicative of reduced anxiety. Moreover, HOM T286A-KI mice also spent more time in the center compared to WT and HET littermates (**Fig 4F**). These data suggest that there is a complex time-dependent relationship between the reductions of CaMKIIα activity and/or expression in these three mouse lines and anxiety in a novel environment.

**Figure 4.**
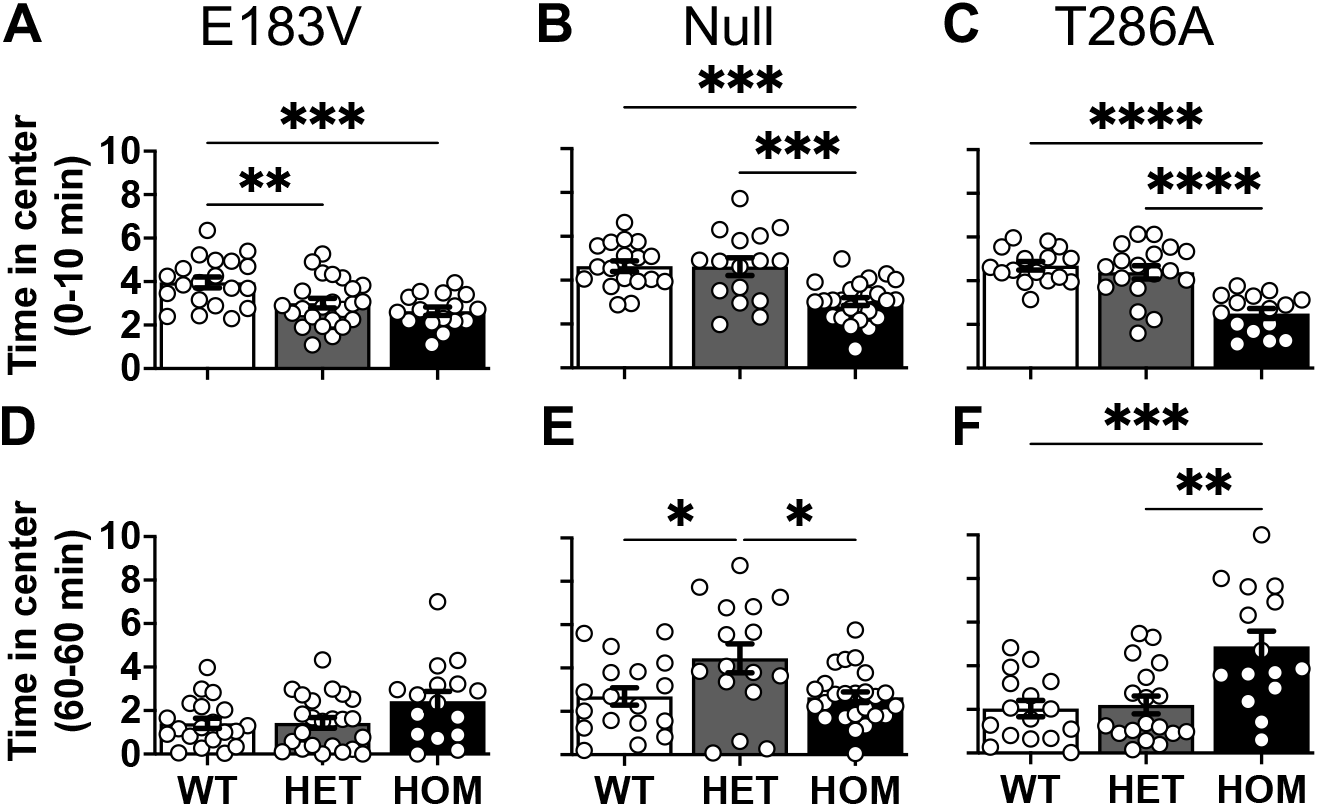
Center time during open-field testing of the three *Camk2a* mutant mouse lines. **A-C:** Time in center during the first 10 mins (mean ± SEM). E183V-KI (**A**): 1-way ANOVA, P=0.0006; Bonferroni’s *post hoc* analysis. Null (**B**): 1-way ANOVA, P<0.0001; Bonferroni’s *post hoc* analysis. T286A-KI (**C**): 1-way ANOVA, P<0.0001; Bonferroni’s *post hoc* analysis. **D-F:** Time in center during the last 10 mins (mean ± SEM). E183V-KI (**D**): 1-way ANOVA, P=0.0431; Bonferroni’s *post hoc* analysis Null (**E**): 1-way ANOVA, P=0.0074; Bonferroni’s *post hoc* analysis. T286A-KI (**F**): 1-way ANOVA, P=0.0003; Bonferroni’s *post hoc* analysis. Symbols indicate P≤0.05 for *, P≤0.01 for **, P≤0.001 for ***and P≤0.0001 for ****. Numbers of WT, HET and HOM mice, respectively, were: E183V-KI: 20, 24, 16. Null: 18, 16, 25. T286A-KI: 16, 18, 15.

To provide more insight into the relationship between *Camk2a* and anxiety, mice were tested for 10 min using an elevated plus maze (EPM). HOM E183V-KI, Null and T286A-KI mice spent significantly more time in the open arms of the maze than their WT or HET littermates, with a corresponding significant decrease in time spent in the closed arms (**Fig 5A-C**). In addition, HOM E183V-KI, Null and T286A-KI made significantly more entries into the open arm of the maze compared to their WT or HET littermates (**Fig 5D-F)**. Moreover, HOM E183V-KI, Null and T286A-KI travelled a greater distance in the open arm of the maze compared to their WT or HET littermates (**Fig 5G-I**). Taken together, the EPM data indicate that HOM CaMKIIα mutant mice from all three lines have reduced anxiety, in contrast to their apparent anxiety during the first 10 min of the open field test.

**Figure 5.**
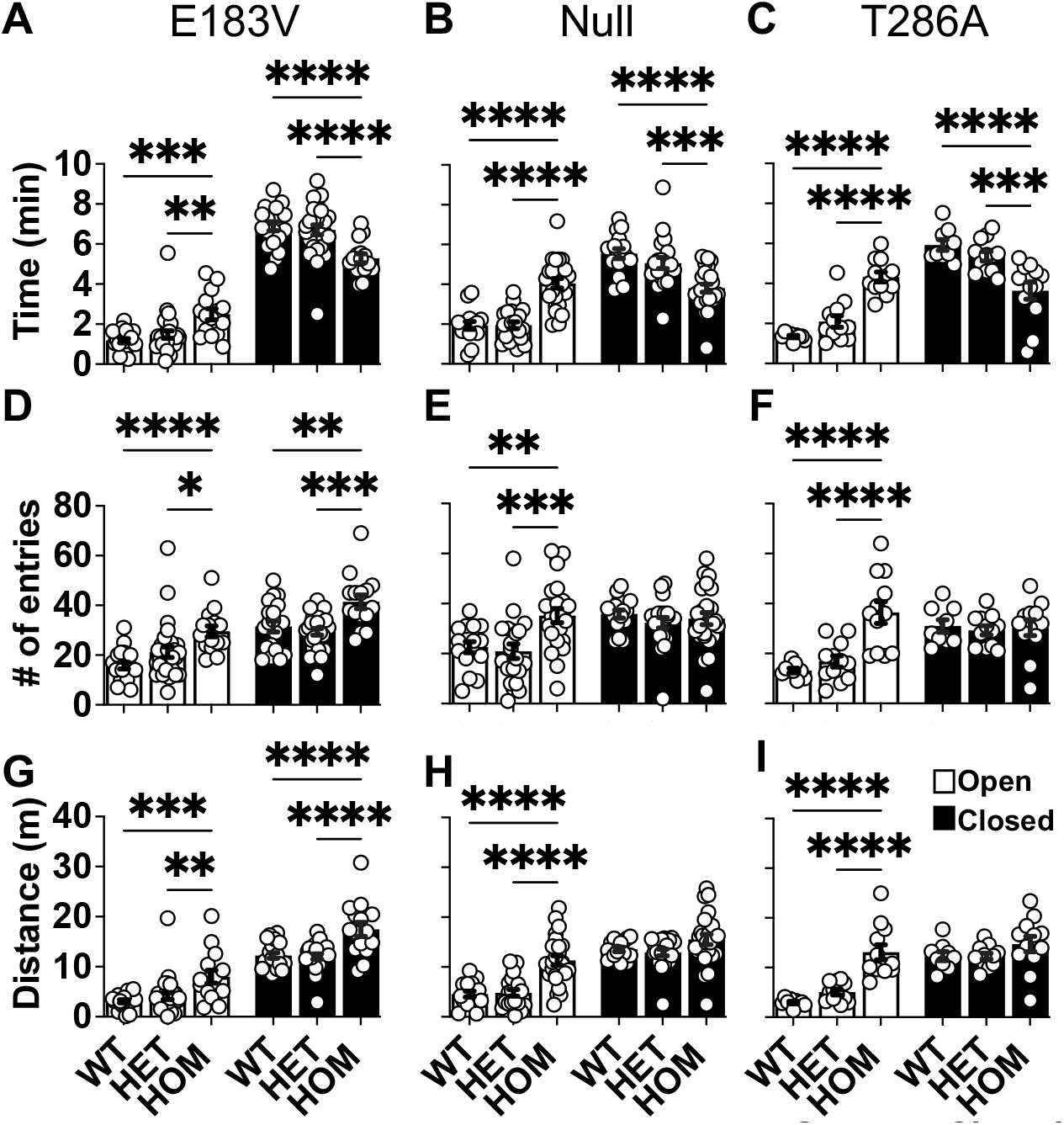
Anxiety-related behaviors during EPM testing of the three *Camk2a* mutant mouse lines. **A-C**: Times spent (mean ± SEM) on the open and closed arms (open and closed bars, respectively) of the EPM for CaMKIIα E183V-KI (**A**), Null (**B**), and T286A-KI (**C**) mice. All data analyzed using a 2-way repeated-measures ANOVA with Tukey’s *post hoc analysis*. E183V-KI (**A**): interaction, F (2, 60) = 11.04, P<0.0001;genotype, F (2, 60) = 2.472, P=0.0930; arm, F (1, 60) = 331.5,P<0.0001. Null (**B**): interaction, F (2, 59) = 24.77, P<0.0001; genotype, F (2, 59) = 8.685, P=0.0005; arm, F (1, 59) = 71.87, P<0.0001. T286A-KI (**C**):interaction, F (2, 30) = 34.51, P<0.0001; genotype, F (2, 30) = 1.021, P=0.3723; arm, F (1, 30) = 77.38, P<0.0001. **D-F**: Numbers of entries (mean ± SEM) onto the open and closed arms of the EPM for CaMKIIα E183V-KI (**D**), Null (**E**), and T286A-KI (**F**) mice. E183V-KI (**D**): interaction, F (2, 60) = 2.860, P=0.0651; genotype, F (2, 60) = 12.66, P<0.0001; arm, F (1, 60) = 72.28, P<0.0001. Null (**E**): interaction, F (2, 59) = 9.900, P=0.0002; genotype, F (2, 59) = 3.954, P=0.0245; arm, F (1, 59) = 26.77, P<0.0001. T286A-KI (**F**): interaction, F (2, 30) = 20.03, P<0.0001; genotype, F (2, 30) = 6.382, P=0.0049; arm, F (1, 30) = 24.82, P<0.0001. **G-I**: Distances traveled (mean ± SEM) on the open and closed arms of the EPM for CaMKIIα E183V-KI (**G**), Null (**H**), and T286A-KI (**I**) mice. E183V-KI (**G**): interaction, F (2, 60) = 1.008, P=0.3710; genotype, F (2, 60) = 15.99, P<0.0001; arm, F (1, 60) = 299.4,P<0.0001. Null (**H**): interaction, F (2, 59) = 6.944, P=0.0020; genotype, F (2, 59) = 15.79, P<0.0001; arm, F (1, 59) = 150.6, P<0.0001. T286A-KI (**I**): interaction, F (2, 30) = 10.87, P=0.0003; genotype, F (2, 30) = 14.31, P<0.0001; arm, F (1, 30) = 75.54, P<0.0001. Numbers of WT, HET and HOM mice, respectively, were: E183V-KI: 20, 29, 15. Null: 16, 21, 25. T286A-KI: 9, 12, 12.

A light-dark box test was then used to further explore the relationship between *Camk2a* and anxiety phenotypes, again with a 10 min test period. There were no significant differences in the amount of time that WT, HET and HOM E183V-KI or T286A-KI mice spent in the light or dark areas of the box (**Figs 6A, 6C**). However, HOM Null mice spent slightly, but significantly, more time on the light side compared to WT mice, indicative of mildly reduced anxiety (**Fig 6B**). Moreover, we found no difference in the number of transitions between the light and dark side within any of the CaMKIIα mouse lines (**Fig 6D-F**). In addition, HOM E183V-KI traveled a significantly greater distance in both the light and dark sections of the box than their littermates (**Fig 6G**).

**Figure 6.**
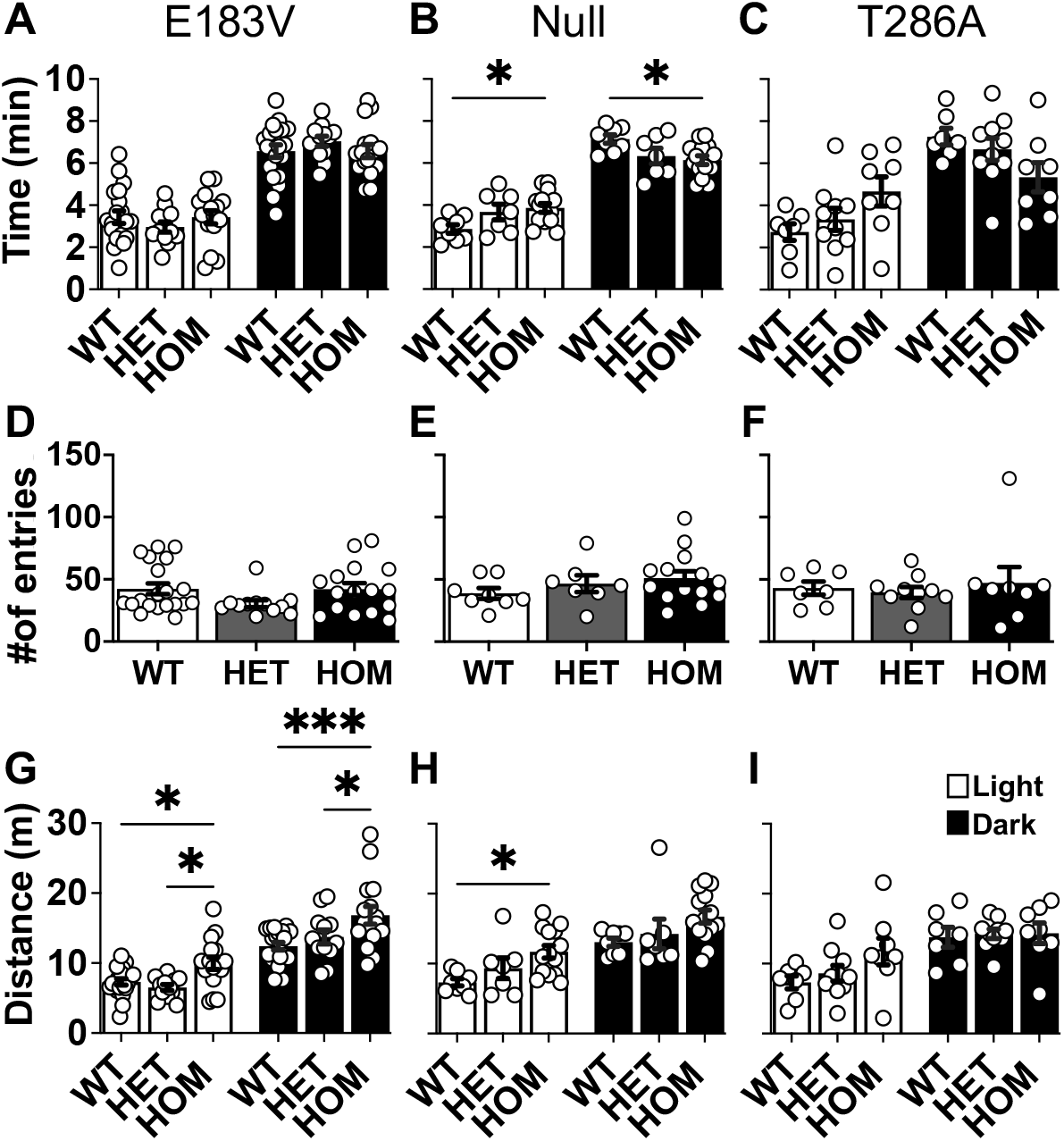
Anxiety-related behaviors during light-dark (LD) testing of the three *Camk2a* mutant mouse lines. **A-C:** Times spent (mean ± SEM) on the light or dark sides (open and filled bars, respectively) of the LD box for CaMKIIα E183V-KI (**A**), Null (**B**), and T286A-KI (**C**) mice. Panels **A-C** and **G-I** analyzed using a 2-way repeated-measures ANOVA with Tukey’s *post hoc analysis* E183V-KI (**A**): interaction, F (2, 45) = 0.6832, P=0.5101; genotype, F (2, 45) = 505.8, P<0.0001; chamber, F (1, 45) = 94.79, P<0.0001. Null (**B**): interaction, F (2, 26) = 14.66, P<0.0001; genotype, F (2, 26) = 0.000, P>0.9999; chamber, F (1, 26) = 349.8, P<0.0001. T286A-KI (**C**): interaction, F (2, 22) = 2.833, P=0.0804; genotype, F (2, 22) = 8.827, P=0.0015; chamber, F (1, 22) = 18.66, P=0.0003. **D-F:** The numbers of entries (mean ± SEM) into the light or dark sides during the LD test for CaMKIIα E183V-KI (**D**), Null (**E**), and T286A-KI (**F**) mice. E183V-KI (**D**): Kruskal-Wallis, P = 0.1712. Null (**E**): 1-way ANOVA, F (2, 26) = 1.2, P=0.3185. T286A-KI (**F**): Kruskal-Wallis, P = 0.8710. **G-I:** Distance traveled (mean ± SEM) on the light or dark sides of the LD box for CaMKIIα E183V-KI (**G**), Null (**H**), and T286A-KI (**I**) mice. All data analyzed using a 2-way repeated-measures ANOVA with Tukey’s *post hoc analysis*. E183V-KI (**G**):interaction, F (2, 45) = 1.341, P=0.2718; genotype, F (2, 45) = 10.22, P=0.0002; chamber, F (1, 45) = 115.3, P<0.0001. Null (**H**): interaction, F (2, 26) = 0.1718, P=0.8431; genotype, F (2, 26) = 4.540, P=0.0204; chamber, F (1, 26) = 75.90, P<0.000. T286A-KI (**I**): interaction, F (2, 22) = 1.136, P=0.3393; genotype, F (2, 22) = 1.813, P=0.1868; chamber, F (1, 22) = 20.84, P=0.0002. Numbers of WT, HET and HOM mice, respectively, were: E183V-KI: 20, 12, 16. Null: 8, 7, 14. T286A-KI, 7, 10, 8.

HOM Null mice also traveled a significantly greater distance in the light section of the box than their WT littermates. In contrast, there were no significant differences in the distance traveled by any genotype of the T286A-KI mice (**Fig 6H-I**). Taken together, the light-dark box data indicate that the complete loss of CaMKIIα activity and expression in CaMKIIα Null mice results in a modest decrease of anxiety-related behaviors, but this is not phenocopied in E183V-KI or T286A-KI mice.

### Tactile phenotypes

Since several ASD mouse models exhibit abnormal tactile responses ^37,38^, we used the von Frey filament test to assess tactile responsiveness of our CaMKIIα mutant mouse lines. More force (thicker filaments) was required to elicit paw withdrawal responses in HOM E183V-KI, Null and T286A-KI mice compared to WT and HET littermates (**Fig 7A-C**). This data show that loss of CaMKIIα activity or expression is sufficient to blunt responses to tactile input.

**Figure 7.**
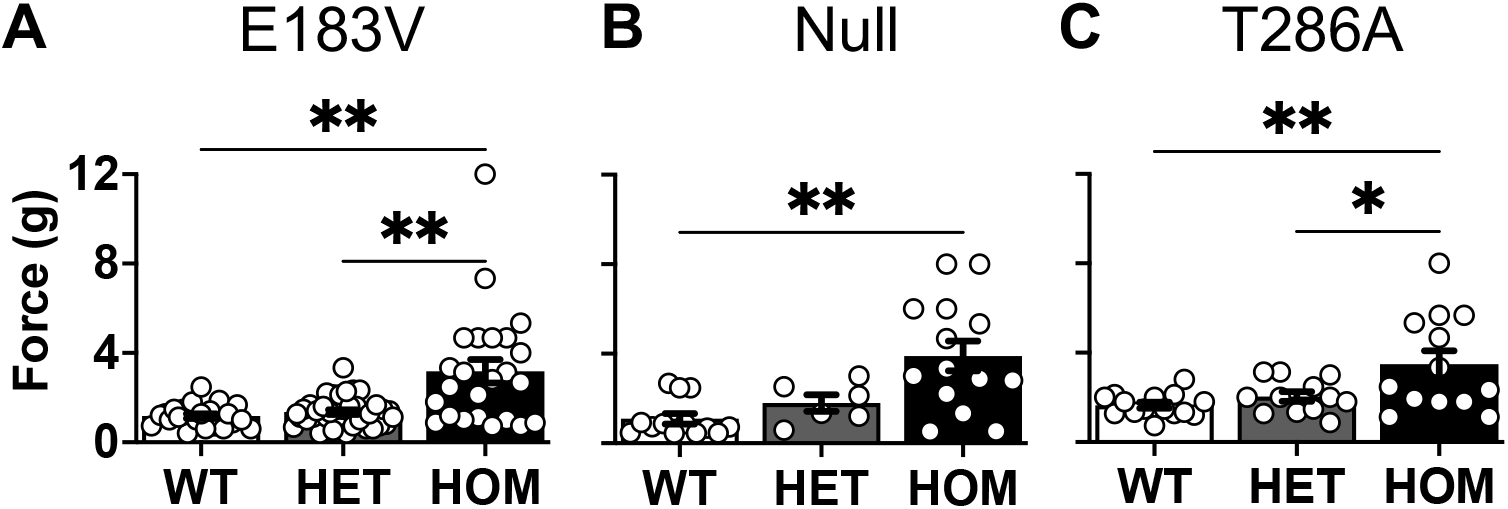
Tactile sensitivity of the three *Camk2a* mutant mouse lines in the VFF test. **A-C:** Force required to evoke paw withdrawal (mean ± SEM). E183V-KI (**A**): Kruskal-Wallis, P=0.0010 with Dunn’s *post hoc* analysis Null (**B**): Kruskal-Wallis, P=0.0021 with Dunn’s *post hoc* analysis T286A-KI (**C**): 1-way ANOVA, P=0.0026 with Bonferroni’s *post hoc* analysis Numbers of WT, HET and HOM mice, respectively, were: E183V-KI: 22, 38, 24. Null: 13, 6, 14. T286A-KI: 15, 12, 13.

## Discussion

In this study, we compared the impact of different loss of function *Camk2a* mutations on mouse behavioral phenotypes often considered to be related to human ASD symptoms ^36,39^. First, we found that reductions in CaMKIIα expression or activity are sufficient to increase stereotypic jumping behavior and hyperactivity. Second, we found that a reduction in CaMKIIα activity and expression impact social exploration with only mild effects on anxiety-related behaviors. Finally, we found that decreases in CaMKIIα expression or activity reduce sensitivity to peripheral tactile input. Taken together, these results show that normal CaMKIIα expression and/or activity is essential for normal motor, social, and sensory functions.

While ASD symptoms typically include social interaction deficits and stereotypic behaviors, there is also a variable spectrum of co-morbidities, often including generalized hyperactivity ^39,40^. Several ASD mouse models also exhibit hyperactivity and stereotypic behaviors, especially in novel environments ^41,42^. In the novel open field test, all three *Camk2a* mutant mouse lines exhibit increases in both stereotypic behavior (jumping; **Fig 1**) and overall locomotor activity (**Fig 2**). Interestingly, the severity of these two phenotypes depends to some degree on the severity of the *Camk2a* mutation. The jumping phenotype is more robust and persistent following the complete loss of CaMKIIα protein and activity in HOM CaMKIIα Null mice compared to HOM E183V-KI or T286A-KI mice. Similarly, the increased motor activity of HOM Null mice persists for at least 60 min in the open-field test, whereas the increased motor activity of HOM T286A-KI and E183V-KI mice is only evident during the first ∼30 min of the novel open field test. Thus, a reduction of only CaMKII activity modestly enhances stereotypic jumping and locomotor activity when combined with the stress associated with entering a novel environment, whereas the complete loss of CaMKIIα expression and activity significantly enhances these phenotypes.

Many ASD mouse models display a range of social interaction deficits ^35,36,43^. However, WT, HET and HOM E183V-KI mice exhibited normal social exploratory behavior in the 3CST (**Fig 3**). HOM Null and T286A-KI mice failed to exhibit a significant preference for social interactions, although there was a trend for a social interaction preference in both lines. Taken together, the current data indicate only mild, if any, impairments in social preferences in the three *Camk2a* mutant mouse lines on a B6 strain background. However, our prior studies of E183V-KI mice on a mixed background detected significant social deficits, and female T286A-KI mice, which were also on a mixed background, were reported to exhibit significant social interaction phenotypes ^26,44^. However, analysis of sex revealed any differences for any *Camk2a* mutant mouse (data not shown). The different genetic background of the mice used in these studies may affect the severity of social interaction deficits. Moreover, it is worth noting that other studies have shown that increasing or decreasing CaMKIIα expression alters aggressive behaviors in mice ^45-47^, perhaps also impacting social behaviors. Taken together, these data indicate that CaMKII mutations have relatively subtle effects on social behaviors that may depend on the strain of mice used.

Initial novel open-field testing indicated that HOM mice in all three *Camk2a* mutant lines spent less time in the center of the arena during the first 10 min of the trials (**Fig 4**), which is typically interpreted as an increase in anxiety. In addition, increased jumping behavior of these mice (**Fig 1**) could be interpreted as a sign of increased anxiety (i.e., a need to escape). While increased anxiety is typically associated with reduced exploratory activity ^36^, HOM mice in all three lines exhibited increased overall motor activity (**Fig 2**). Moreover, the novelty-induced stress of the open-field arena may have contributed to the apparent anxiety because the decrease in center time was not detected during the last 10 min (of the 60 min trial). In fact, there was an increased center time for HOM T286A-KI mice during the last 10 min, and also a trend for increased center time with HOM E183V-KI mice, indicative of reduced anxiety.

To provide greater clarity about possible anxiety-related phenotypes, we also tested *Camk2a* mutant mice in the EPM and light-dark box. However, these studies provided conflicting insight into the role of CaMKIIα in anxiety. In the EPM, WT mice, as well as HET littermates from all three lines spent 3-8 times more time in the closed arm vs. the open arm. HOM mice from all three lines spent significantly more time in the high-risk open arms than their WT or HET littermates (**Fig 5**). In fact, HOM Null and T286A-KI mice failed to display any preference for the closed arms (closed/open time ratio 1.1 and 0.9, respectively), and the closed/open time ratio for HOM E183V-KI mice was significantly reduced relative to WT or HET littermates (E183V-KI closed/open arm time ratios: WT, 8.2 ± 1.5; HET, 6.3 ± 1.0; HOM, 2.8 ± 0.5; p= 0.0001by 1-way ANOVA). In contrast to the apparently decreased anxiety of all three HOM mice in the EPM, the light-dark box revealed more modest, if any, anxiety phenotypes. In fact, only HOM Null mice displayed a significant change in preference for the light zone in the box relative to WT littermates, indicative of a mild decrease in anxiety, with no significant effects of the other *Camk2a* mutations on light-dark preference (**Fig 6**). Interestingly, the EPM also revealed significant increases of distances covered exploring high risk area (i.e. open arm) for HOM mice from all three lines, and this increase of high-risk exploration was confirmed for E183V-KI and Null mice in the light-dark box. In fact, before the EPM was complete, some HET and HOM mice would jump off the open arm, resulting in their exclusion from the analyses. A similar phenomenon was observed in pilot studies using an elevated zero maze (data not shown). Taken together, these data indicate that all three of these *Camk2a* mutations increase exploratory and impulsive behaviors, which may be misinterpreted as changes in anxiety in some testing paradigms.

Alterations in various forms of sensory perception are another common co-morbidity of ASD ^48,49^, and several ASD mouse models display altered tactile sensitivity ^37,38^. Previous studies of *Camk2a* Null and T286A-KI mice revealed alterations in the sensitivity to whisker stimulation ^50-52^. Here we used the von Frey filament test to show for the first time that reduced CaMKIIα expression and/or activity in any of the three *Camk2a* mutant mouse lines lead to a substantial loss of tactile sensitivity in the paw (**Fig 7**). Moreover, the sensitivity to a nociceptive electric foot shock was also shown to be increased by the loss of CaMKIIα expression ^45^, and CaMKIIα has been linked to the development of chronic peripheral pain conditions such neuropathic and inflammatory pain ^53-56^. While von Frey filaments do not appear to induce nociceptive responses, it will be interesting to investigate the role of CaMKIIα in responses to more noxious stimuli, such as foot shock or heat.

## Conclusion

The current findings add to the diversity of mouse behavioral phenotypes that result from global genetic disruptions of CaMKIIα activity or expression. This diversity is not surprising given that CaMKIIα is widely expressed in many, but not all, parts of the forebrain, as well as in cerebellar Purkinje cells and spinal cord. Since prior studies have shown that one or more of these *Camk2a* mutation can disrupt normal excitatory synaptic transmission in the hippocampus, cortex, striatum, cerebellum and other brain regions, it seems likely that the diverse behavioral phenotypes observed here and in prior studies result from abnormal glutamate receptor function and synaptic plasticity in one or more brain regions. Further studies are required to reveal the relative contributions of disrupted synaptic function in specific brain regions to the behavioral phenotypes of CaMKIIα mutant mice.

## Supporting information

Complete statistical reports on all data

## Acknowledgments

All behavioral experiments were performed in part through the use of the Vanderbilt Murine Neurobehavior Core lab supported in part by the EKS NICHD of the NIH under Award #U54HD083211. The content is solely the responsibility of the authors and does not necessarily represent the official views of the NIH. This study was funded by: Burroughs Wellcome Fund Postdoctoral Enrichment Program and Postdoctoral Training Program in Functional Neurogenomics (T32-MH065215)

